# The formal demography of kinship IV: Two-sex models

**DOI:** 10.1101/2022.01.17.476606

**Authors:** Hal Caswell

## Abstract

**Background:** Previous kinship models analyze female kin through female lines of descent, neglecting male kin and male lines of descent. Because males and females differ in mortality and fertility, including both sexes in kinship models is an important unsolved problem.

**Objectives:** The objectives are to develop a kinship model including female and male kin through all lines of descent, to explore approximations when full sex-specific rates are unavailable, and to apply the model to several populations as an example.

**Methods:** The kin of a focal individual form an age×sex-classified population and are projected as Focal ages using matrix methods, providing expected age-sex structures for every type of kin at every age of Focal. Initial conditions are based on the distribution of ages at maternity and paternity.

**Results:** The equations for two-sex kinship dynamics are presented. As an example, the model is applied to populations with large (Senegal), medium (Haiti), and small (France) differences between female and male fertility. Results include numbers and sex ratios of kin as Focal ages. An approximation treating female and male rates as identical provides some insight into kin numbers, even when male and female rates are very different.

**Contribution:** Many demographic and sociological parameters (e.g., aspects of health, bereavement, labor force participation) differ markedly between the sexes. This model permits analysis of such parameters in the context of kinship networks. The matrix formulation makes it possible to extend the two-sex analysis to include kin loss, multistate kin demography, and time varying rates.

## 1 Introduction

The kinship models of Caswell (2019a, 2020); Caswell and Song (2021), following those of Goodman, Keyfitz, and Pullum (1974), and including the model of Coste et al. (2021), describe the kinship network of a focal individual using only female demographic rates. The result is a projection of female kin (e.g., daughters, granddaughters, …) through female lines of descent (e.g., granddaughters include daughters of daughters, but not daughters of sons). This paper removes this limitation and presents a complete two-sex version of the matrix analytic kinship model. The kinship network is defined relative to a focal individual, referred to as Focal. The model provides the age and sex structures of all types of kin through all lines of descent. Female and male rates generally differ, and this model makes it possible for the first time to explore the effects of these differences on the kinship network.

Differences between female and male rates are well known. Female longevity almost always exceeds male longevity. The female advantage in life expectancy, on the order of 2–10 years, has long been documented (Dublin, Lotka, and Spiegelman, 1949). In an analysis of all countries, Raftery, Lalic, and Gerland (2014) found the median female advantage, from 1950 to 2010, to range from 2.5 to 6 years, with some values as high as 10–12. Clark and Peck (2012) found life expectancy gaps of 3–8 years in the late 20th century in an analysis of 195 countries, and related the gap to factors including women’s status, traditional male hazards, development, income inequality, and female representation in government. Glei and Horiuchi (2007), analyzing 29 high income countries, found the gap affected by changes not only in the levels but also in the age patterns of male and female mortality. The gaps in life expectancy are also reflected in differences between women and men in healthy longevity and the proportion of life spent in poor health (Luy and Minagawa, 2014; Oksuzyan et al., 2014).

Male and female fertility can also differ substantially. The most obvious difference is that men can reproduce at much later ages than women (Kühnert and Nieschlag, 2004; Bribiescas, 2016; Paget and Timaeus, 1994). This difference has been used to great effect by Tuljapurkar, Puleston, and Gurven (2007) as a compelling explanation for the evolution of post-reproductive survival in human females.

In addition to differences in timing, the levels of male and female reproduction often differ.^1^ In a valuable recent review, Schoumaker (2019) analyzes male and female fertility, ages at parenthood, and trends in these quantities for 160 countries, and provides an extensive source of data in the online appendices to the article (Schoumaker, 2019, Appendix A). He reports that male TFR (total fertility rate) almost always exceeds female TFR, by an amount that increases with female TFR. Thus as a population goes through the demographic transition, female and male fertility become more similar. The mean age at paternity (the mean age of childbearing for men) is always greater than the mean age at maternity (mean age of childbearing for women), by five to fifteen years. Schoumaker presents the age-specific fertility rates for men and women in Senegal, Haiti, and France (his Figure 4; these will be used below in Section 4), as typical results for high, medium, and low fertility populations and explores the roles of overall fertility, economics, and polygyny in determining these patterns.

Incorporating the demographic rates of both sexes makes it possible to explore the consequences of the differences between female and male rates. In particular, questions about the sex composition of various kinds of kin, as a function of the age of Focal, can be addressed in this new framework. It is known that differential mortality leads to a female-skewed population structure among the elderly (e.g., 10.8 females per male among supercentenarians; Robine and Vaupel 2001). We can ask how this skew will differ among the various kinds of kin,

The attentive reader is no doubt aware that complete sets of male and female rates are not exactly common, which is true. With this in mind, the model presented here accepts any level of sex-specificity in the data: (a) age-specific mortality and fertility schedules for both women and men, (b) sex-specificity in only mortality or only fertility, or (c) no sex-specific data at all, treating the sexes as identical. The last option does away with all the interesting effects of the differences between women and men, but still allows an approximate accounting for the numbers of male and female kin through male and female lines of descent.

The two-sex kinship model includes both males and females in the population state vector, much as Caswell (2019a) incorporated living and dead kin and Caswell (2020) incorporated age and parity status. Projection matrices that incorporate both male and female rates are used to generate the dynamics of all types of kin.

The model presented here is linear. In principle, two-sex fertility rates in sexually reproducing species must depend on the relative abundance of males and females. This leads to frequency-dependent nonlinearities embodied in a marriage or mating function (e.g., Keyfitz, 1972; Iannelli, Martcheva, and Milner, 2005; Shyu and Caswell, 2018). However, this effect would operate not within the population of a particular type of kin, but in the population as a whole, which would require an additional model, not considered here. This model, as with most demographic analyses, is conditional on the hypothesis that male and female fertility schedules remain in effect throughout the calculation. The incorporation of a fully nonlinear two-sex model remains an open research problem.

### 1.1 Notation and terminology

Matrices are denoted by upper case bold characters (e.g., **U**) and vectors by lower case bold characters (e.g., **a**). Female and male rates are distinguished by subscripts (e.g., **U**_f_ and **U**_m_). Vectors are column vectors by default; **x**^⊤^ is the transpose of **x**. The *i*th unit vector (a vector with a 1 in the *i*th location and zeros elsewhere) is **e**_*i*_. The vector **1** is a vector of ones, and the matrix **I** is the identity matrix. When necessary, subscripts are used to denote the size of a vector or matrix; e.g., **I**_*ω*_ is an identity matrix of size *ω* × *ω*. Matrices and vectors with a tilde (e.g., **Ũ** or **ã**) are block-structured, containing blocks for females and males.

The symbol ° denotes the Hadamard, or element-by-element product (implemented by. * in Matlab and by * in R). The symbol ⊗ denotes the Kronecker product. The vec operator stacks the columns of a *m* × *n* matrix into a *mn* × 1 column vector. The notation ‖**x**‖ denotes the 1-norm of **x**. On occasion, Matlab notation will be used to refer to rows and columns; e.g., **F**(*i*, :) and **F**(:, *j*) refer to the *i*th row and *j*th column of the matrix **F**.

## 2 Principles of the two-sex kinship model

Before presenting the complete derivation (Section 3), it is helpful to consider the principles along which the model is organized. The kinship network, as shown in Figure 1, consists of descendants of Focal (children, grandchildren, etc.), ancestors of Focal (parents, grandparents), and descendants of ancestors (siblings, aunts, cousins, etc.). Each kin type is identified by a letter; these become variables in the model.

**Figure 1:**
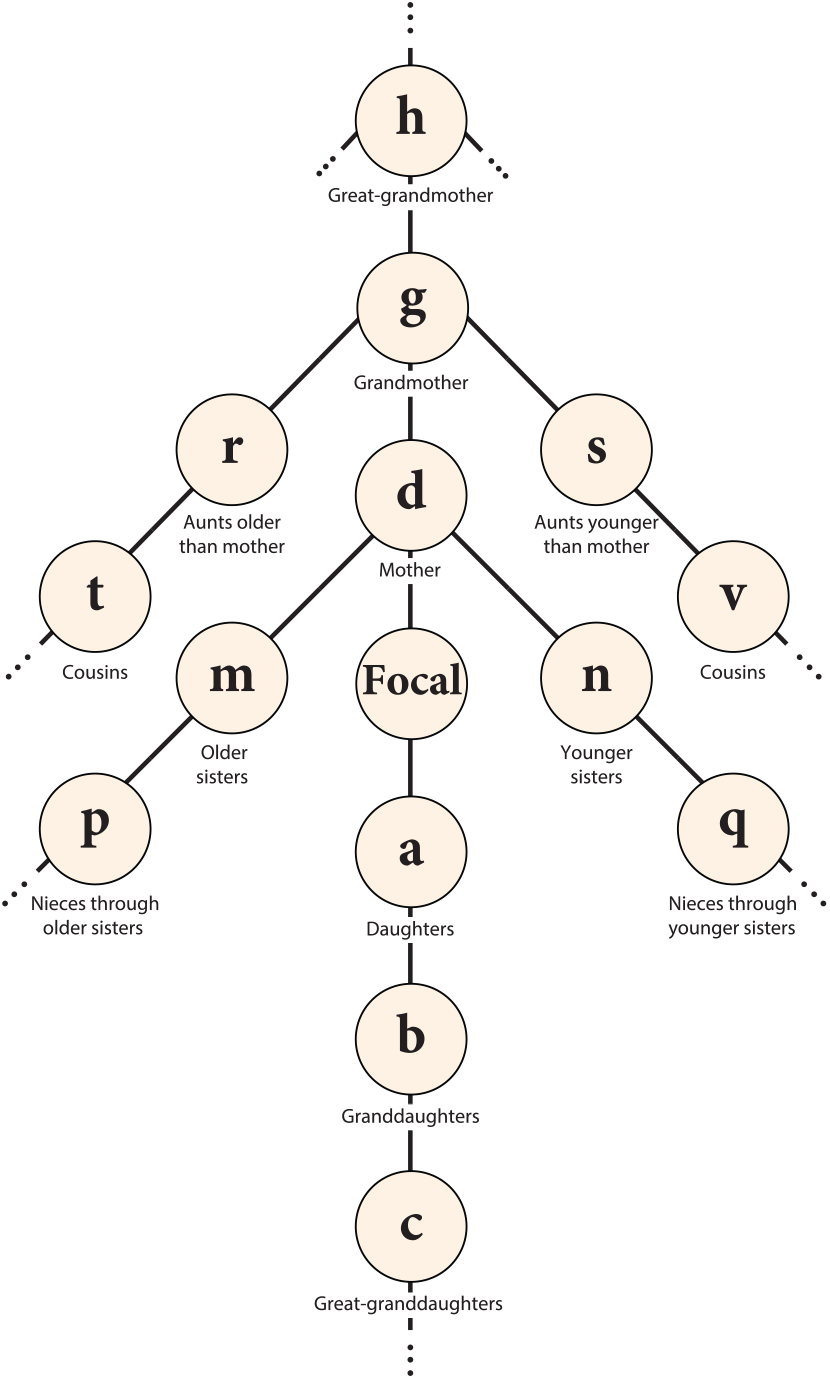
The kinship network surrounding the Focal individual. Symbols (**a, b**, etc.) denote the age structure vectors of each type of kin of Focal. Modified from Caswell (2019a) based on network defined in Goodman, Keyfitz, and Pullum (1974) and Keyfitz and Caswell (2005).

Each kin type contains a set of subtypes defined by sex and line of descent. These will be compressed, under a reasonable set of assumptions, to individuals classified by age and sex. To show the principles, consider the descendants and the ancestors of Focal.

### 2.1 Descendants of Focal

In Figure 2, Focal produces two types of children, daughters and sons. These give rise to four types of grandchildren: daughters of daughters, daughters of sons, sons of daughters, and sons of sons. This chain of descendants extends naturally to eight types of great-grandchildren, and so on.

**Figure 2:**
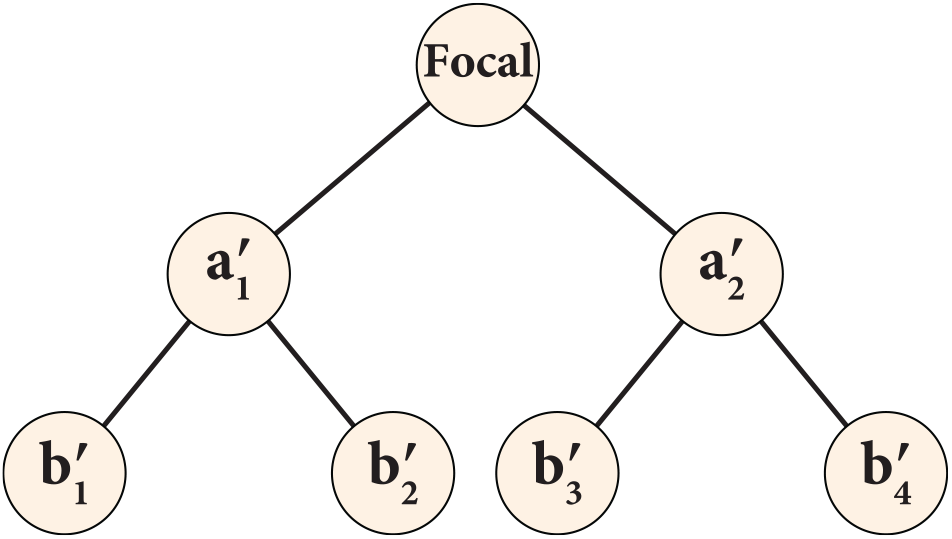
Female and male children (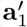 and 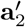) and grandchildren 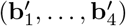 of Focal. The symbols represent age distribution vectors; the primes indicate that kin are defined by sex and by line of descent.

The vectors 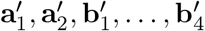 are the age structure vectors of each of these types of kin. Our convention is to number females first (1, 3, 5, …) and males second (2, 4, 6, …). The primes appearing in Figure 2 indicate that those vectors are specific to both sex and line of descent. (We will eliminate them shortly.)

We begin by specifying notation. Define

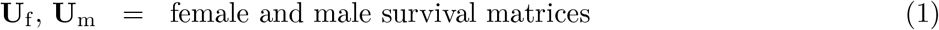

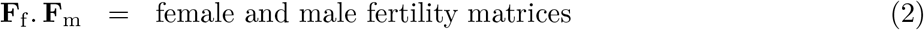

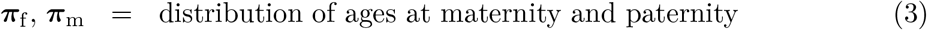

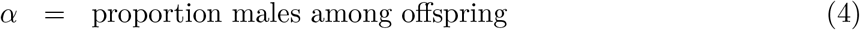

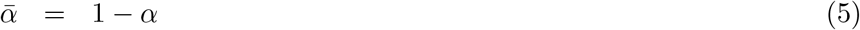

survival matrices contain age-specific survival probabilities on the subdiagonal and zeros elsewhere. Fertility matrices contain age-specific fertility rates on the first row and zeros elsewhere. The proportion of males at birth *α* = 0.5 throughout. The distributions ***π***_f_ and ***π***_m_ are defined in Section 3.1.

For the case of the children in Figure 2, we write

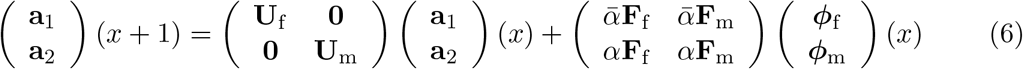

Daughters and sons survive according to the matrices **U**_f_ and **U**_m_. New children are produced by the fertility of Focal. If Focal is a female, then ***ϕ***_f_(*x*) = **e**_*x*_, where **e**_*x*_ is a vector of length *ω* with a 1 in the *x*th entry and zeros elsewhere, and ***ϕ***_m_(*x*) = **0**. If Focal is male, then these vectors are reversed.

The dynamics of grandchildren must account for all four types of grandchildren shown in Figure 2:

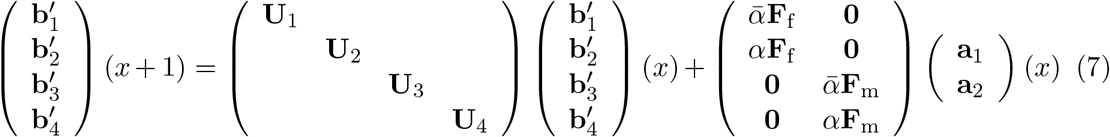

As written, equation (7) permits each of the four types of grandchildren to experience its own survival schedule, and each of the two types of children to contribute according to its own fertility schedule. The corresponding model for great-grandchildren would include eight survival matrices, **U**_1_, …, **U**_8_ and four fertility matrices **F**_1_, …, **F**_4_. And so on.

Let us make the usual assumption that the demographic rates are affected by sex but not by line of descent from an arbitrarily defined Focal individual. Then

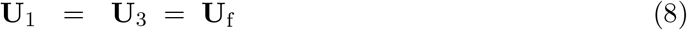

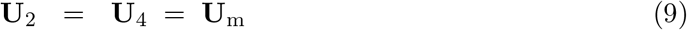

Under this assumption, the four types of grandchildren can be aggregated into granddaughters and grandsons, ignoring their lines of descent from Focal. Define

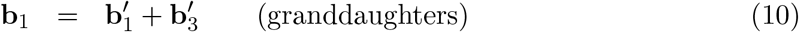

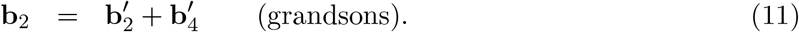

The dynamics then become

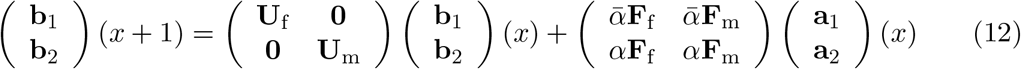

This pattern continues for subsequent generations of descendants, leading to the pattern shown in Figure 3, in which female and male children, grandchildren, and so on are produced by the female and male survival and fertility matrices.^2^

**Figure 3:**
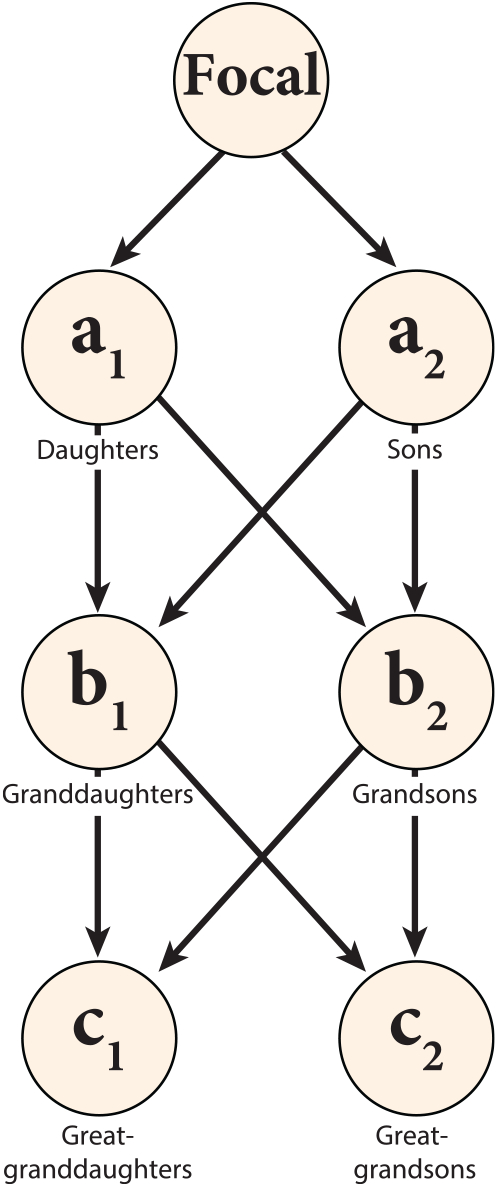
Female and male children, grandchildren, and great-grandchildren of Focal. Obtained from the graph in Figure 2 by combining male and female kin, regardless of lines of descent.

A final notational simplification follows from writing the vectors containing female and male age distributions as

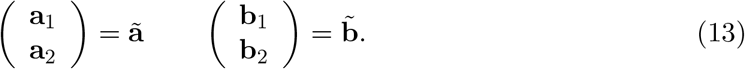

If appropriate block-structured matrices are defined, Figure 3 reduces to Figure 4 and equations (6) and (12) simplify to the projections of a single vector. Section 3 develops the entire kinship model in these terms.

**Figure 4:**
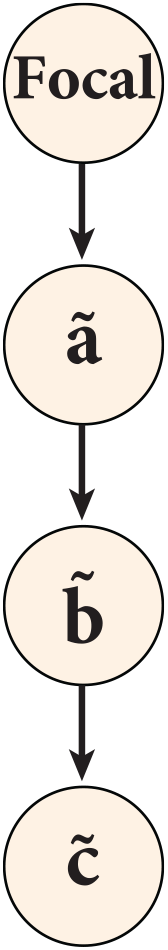
The children, grandchildren, and great-grandchildren of Focal. The female and male kin in Figure 3 have been combined into the block-structured vectors **ã**, 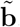, and 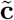.

### 2.2 Accounting for ancestors

Focal has a network of female and male ancestors, as shown in Figure 5. She has two parents (mothers 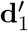 and fathers 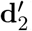), four grandparents (maternal 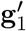 and 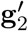; paternal 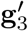 and 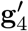). The pattern continues for as many levels as desired, the number of types of ancestors doubling with each level.

**Figure 5:**
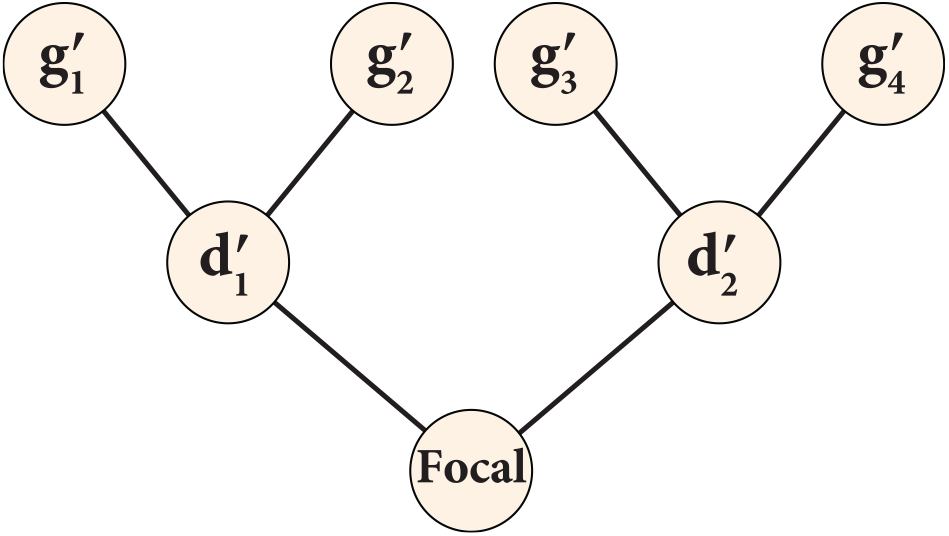
Accounting for the parents (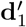 and 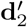) and grandparents 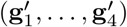 of Focal. Symbols represent age structure vectors; primes indicate that individuals are characterized by both sex and chains of ancestry.

Consider the grandparents. Their dynamics satisfy

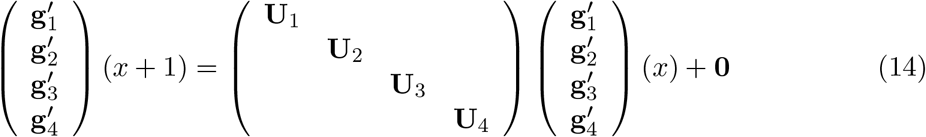

where the recruitment term is zero because Focal accumulates no new biological ancestors after her birth. Combining the maternal and paternal grandparents of each sex yields

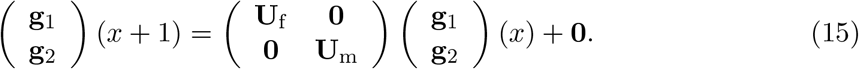

### 2.3 Reproduction by ancestors

The reproduction of the ancestors of Focal produces the side chains in Figure 1. In the two-sex model, reproduction must account for the lack of independence of pairs of ancestors. Consider Focal’s daughters and sons. They can be assumed to reproduce independently to produce grandchildren, and are treated so in Figure 3 and equation (12). But Focal’s mother and father do not reproduce independently to produce her siblings. Therefore, as shown in Figure 6, we credit the reproduction of Focal’s parents to her mother, following the female fertility schedule (a kind of female dominance assumption).

**Figure 6:**
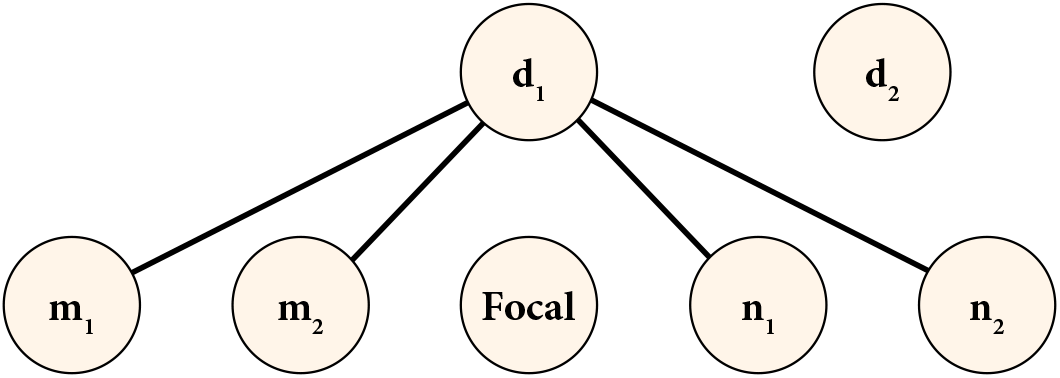
An example of reproduction by ancestors (parents, in this case) of Focal. All reproduction by Focal’s parents, producing older and younger sisters and brothers, is attributed to Focal’s mother.

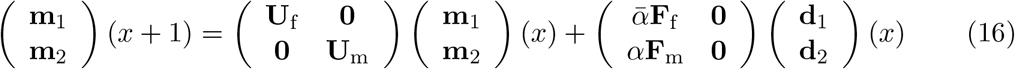

Notice the difference between the block-structured fertility matrices in (16) and (12).

## 3 Two-sex kinship dynamics

For some type **k** of kin, we write

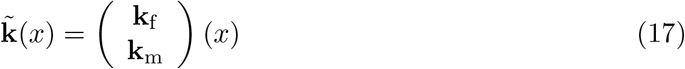

where *x* is the age of Focal. The tilde denotes block structured vectors and matrices composed of female and male parts.

The dynamics of **k**(*x*) are written as

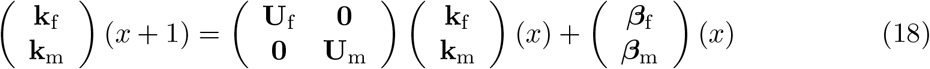

or

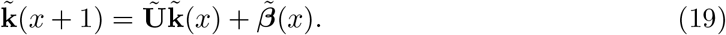

The recruitment subsidy term 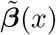 depends on the nature of the kin that provide the recruitment.

- If the subsidy is provided by reproduction of one of the direct ancestors of Focal (parents, grandparents, etc.), then, as in equation (16)

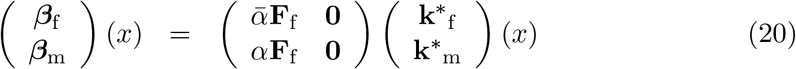

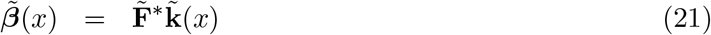

where **k**\* denotes the source kin (e.g., mothers are the source of the siblings of focal). The matrix 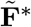 captures reproduction of both female and male offspring by females.
- If the subsidy is provided by any other kin type, then as in equation (12)

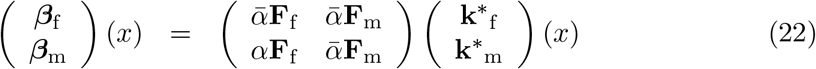

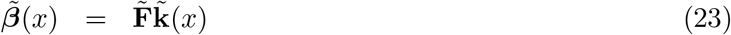

where **k**\* again denotes the source kin (e.g., children are the source of the grandchildren of focal).
- If there is no recruitment subsidy, as in (15), then 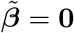.

### 3.1 Ages at maternity and paternity

The matrices **F**_f_ and **F**_m_ contain age-specific fertilities for females and males respectively. The distributions of the ages of mothers and of fathers are obtained by applying these per capita rates to age distributions of women and men, respectively. Let

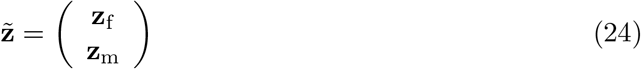

be the age structure of the population. The age distributions of maternity and paternity are

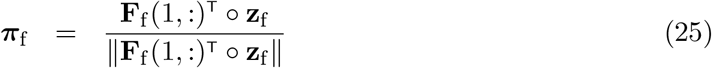

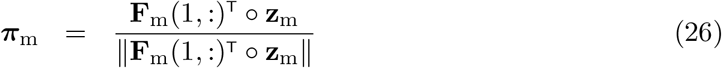

and we write

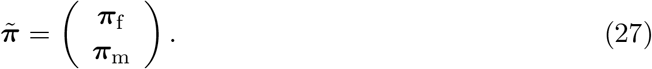

The age structure 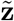 could be obtained from projection of a previous population (Caswell and Song, 2021) or from an observed population structure. Here, following Goodman, Keyfitz, and Pullum 1974; Caswell 2019a, 2020, we use the stable population defined by the demographic rates in **Ũ** and 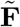. Define a projection matrix for the female dominant population

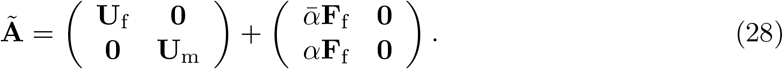

The stable sex-age structure is given by the eigenvector 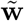 corresponding to the dominant eigenvalue of **Ã**. Without loss of generality, we scale 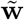 so that its entries are non-negative, write

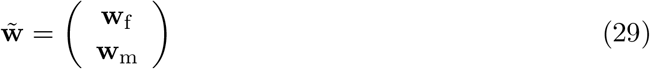

and substitute 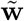 for 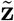 in (24). Notice the use of 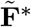 rather than 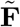; this corresponds to the usual female dominance assumption in stable population theory.

### 3.2 Kinship dynamics

We turn now to the model for each type of kin.

#### 3.2.1 Focal

Focal is an individual of specified sex, alive at age *x*. The state of Focal is given by the vector

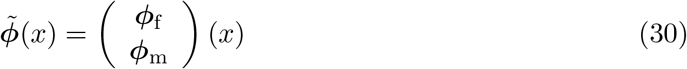

This can be extended to cases where Focal is classified by stage as well as age, following the methods in Caswell (2020).

#### 3.2.2 Children and descendants of Focal

The dynamics of the children of focal are given by

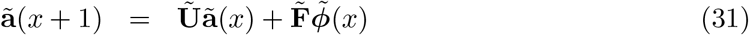

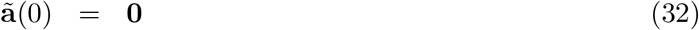

The dynamics of the grandchildren of focal are given by

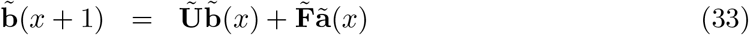

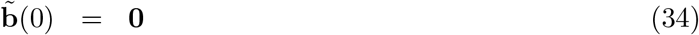

The dynamics of the great-grandchildren of focal are given by

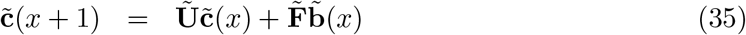

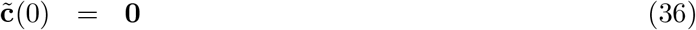

In each case, the initial condition is zero (Focal has no children, grandchildren, etc. when she is born). The recruitment of each generation of descendants comes from the fertility of the previous generation.

The chain of descendants can be extended as far as desired, as is also true of the matrix models of Caswell (2019a, 2020); Caswell and Song (2021), To do so, let 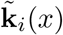 be the state vector of the *i*th generation of descendants, with Focal defined as generation 0. Then

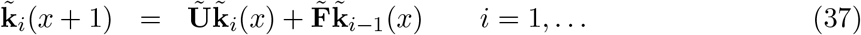

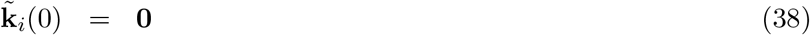

#### 3.2.3 Parents and ancestors of Focal

Counting the ancestors (parents, grandparents, etc.) of Focal involves going up the branching network shown in Figure 5. We know that, at birth, Focal had exactly one living mother. We will assume that she also has one living father, thus ignoring paternal mortality in the nine months between conception and birth.

The ages of Focal’s parents at her birth are unknown, so we treat her mother and father as being selected at random from the distributions ***π***_f_ and ***π***_m_ of the ages of mothers and fathers at the birth of children.

Under these assumptions, the dynamics of the parents of Focal are given by

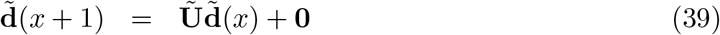

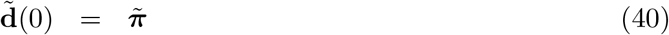

Focal accumulates no new parents after her birth, so the recruitment term is zero.

The dynamics of the grandparents of Focal are given by

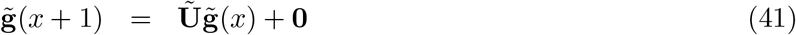

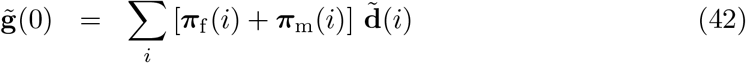

The initial condition 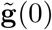 is obtained by noting that the grandparents of Focal are the parents of Focal’s parents. Thus we could write

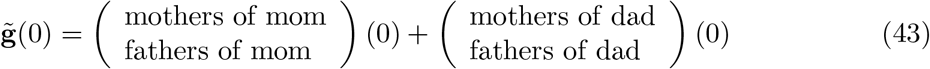

We do not know the ages of Focal’s mother or father, but we know their distributions, so

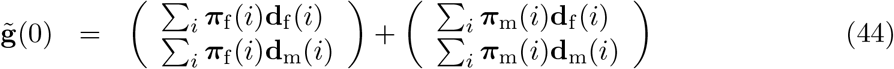

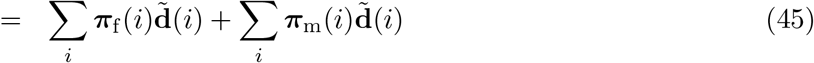

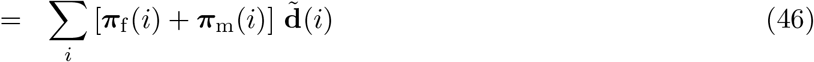

The dynamics of the great-grandparents of Focal follow the same pattern. The great-grandparents of Focal at her birth are the grandparents of Focal’s parents, so

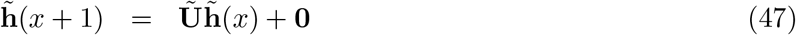

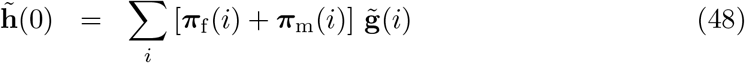

As with descendants, the ancestors can be calculated back as far as desired. For this calculation, let 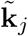 be the kin vector of the *j*th generation of ancestors, where parents are generation zero. Then, for *j* ≥ 1,

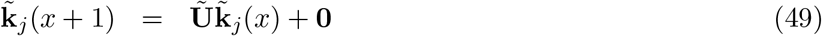

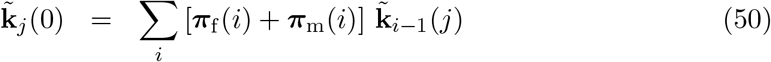

#### 3.2.4 Siblings, nieces, and nephews of Focal

We turn next to the siblings, nieces, and nephews of Focal. Inconveniently, English seems to have no gender neutral collective term for nieces and nephews, as sibling is for brothers and sisters. The term “nibling” has been suggested, according to the internet. German has such a term, “Geschwisterkind,” meaning child of a sibling. There also seems to be no gender-neutral collective term for aunts and uncles (next section).

##### Siblings

The older and younger siblings of Focal are treated separately because they have different dynamics.

Focal may have older siblings at her birth, but she can accumulate no more of them after she is born, so the recruitment term is zero. Older siblings at Focal’s birth are the children of Focal’s mother at Focal’s birth. Focal’s mother and father do not reproduce independently, so the initial condition is credited to Focal’s mother and calculated as an average over ***π***_f_ only. Thus

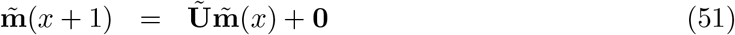

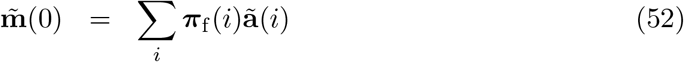

Focal has no younger siblings at her birth,^3^ but can accumulate younger siblings through her mother’s reproduction. Thus

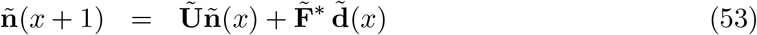

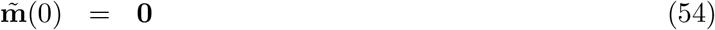

The matrix 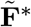 ensures that the recruitment of new younger siblings comes from the reproduction of Focal’s mother following the female fertility schedule, as in Figure 6.

##### Niblings

Then niblings of Focal are the children of Focal’s older and younger siblings.

The recruitment of niblings through older siblings comes from the reproduction of those siblings, both brothers and sisters contributing independently. The initial condition follows from the fact that, at the time of Focal’s birth, these niblings are the grandchildren of Focal’s mother. Thus, the dynamics of nieces and nephews through older siblings are

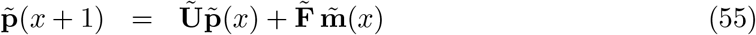

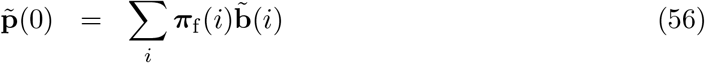

This line of descent can be continued indefinitely. Each generation receives its recruitment from the generation before (grand-niblings from niblings, etc.). The initial condition for each generation is the corresponding descendant of Focal’s mother: niblings are the grandchildren of Focal’s mother; grand-niblings are the great-grandchildren, and so on.

The recruitment of niblings through younger siblings comes from the reproduction of those younger siblings. The initial condition is zero, because Focal has no younger siblings at birth, and hence can have no nieces or nephews through them. The resulting dynamics are

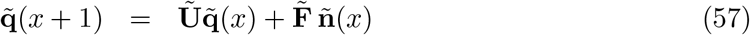

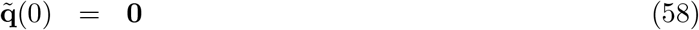

#### 3.2.5 Aunts and uncles of Focal

The aunts and uncles of Focal (there seems to be no gender-inclusive term) are the siblings of Focal’s parents.

The aunts and uncles through the older siblings receive no recruitment subsidy because once Focal is born, her parents cannot add any older siblings. The initial condition combines the older siblings of the mother and father of Focal, following the steps above for the grandparents of focal. The resulting dynamics are

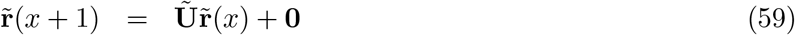

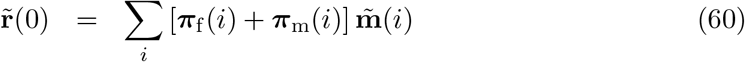

Aunts and uncles through younger siblings receive a recruitment subsidy from the reproduction of the grandmothers of Focal. Grandmothers and grandfathers do not reproduce independently, so only input from grandmothers is counted. The initial condition combines the younger siblings of Focal’s mother and of Focal’s father, at the time of Focal’s birth.

The resulting dynamics are

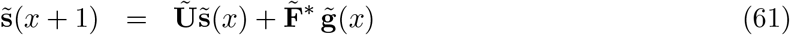

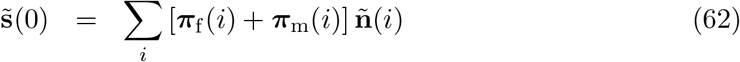

#### 3.2.6 Cousins of Focal

The cousins of Focal are the children of the aunts and uncles of Focal. The cousins through the older aunts/uncles receive a recruitment subsidy from the reproduction of older aunts and uncles. These cousins are the nieces and nephews of Focal’s mother through her older siblings.

The resulting dynamics are

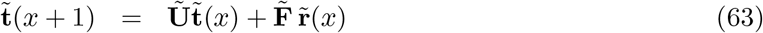

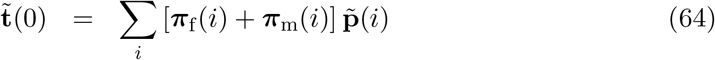

The cousins through the aunts/uncles younger than mother receive a recruitment subsidy from the reproduction of those aunts and uncles. These cousins are the nieces and nephews of Focal’s mother through her younger siblings.

The resulting dynamics are

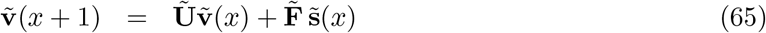

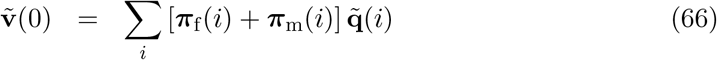

The chains of descendants through these cousins can be extended indefinitely. Recruitment into each generation comes from the generation before it, and the initial condition consists of the corresponding level of nieces/nephews of Focal’s mother.

## 4 Some two-sex kinship patterns: Senegal, Haiti, France

As an example, I explore some two-sex kinship patterns using mortality and fertility schedules for Senegal (2013), Haiti (2010), and France (2012). These were used by Schoumaker (2019) as examples of populations with large, medium, and small differences between female and male fertility schedules; see Table 2. In Senegal, male TFR is much greater than female TFR and mean age at paternity is much greater than mean age at maternity. In France, the differences are reduced even further. The three populations also differ in mortality, and all three show typical differences between female and male life expectancy.

### 4.1 Fertility schedules

Age-specific fertility schedules in five-year age classes were kindly provided by Bruno Schoumaker based on information in Schoumaker (2019). The five-year age classes were interpolated to one-year age intervals, using the Matlab function `interp1` with cubic spline interpolation (Matlab method `akemi`). The observed fertility rates and the interpolated rates used in the model are shown in Figure 7.

**Figure 7:**
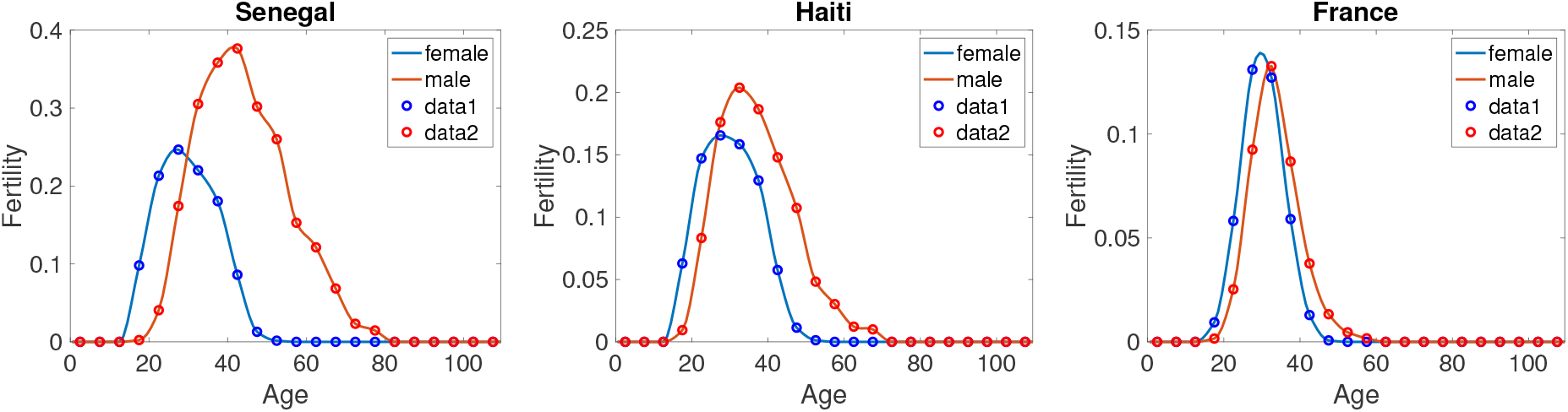
The observed (circles) and interpolated (lines) age-specific fertility rates for Senegal, Haiti, and France. Based on data from Schoumaker (2019).

### 4.2 Mortality schedules

Mortality data for the three countries were obtained from United Nations compilations^4^ (United Nations, 2019). Survival probabilities were taken from abridged male and female life tables for the time period 2010–2015. These five-year survival probabilities were transformed to one-year probabilities, and then interpolated to one-year age intervals using cubic spline interpolation. The UN life tables are truncated at 85 years of age; I extended them to age 100 by assuming that survival declined to 0 at age 100 and extrapolating the values between age 85 and 100 using cubic splines.

### 4.3 Numbers of female and male kin

Of the many results that could be calculated from the model, here I show results for the numbers of female and male kin and sex ratios for each kind of kin, as a function of the age of Focal. Results for kin numbers are shown in Figure 8. The three vertical columns represent, from left to right, Senegal, Haiti, and France. Each row is a different type of kin.

**Figure 8:(part 1 of 2).**
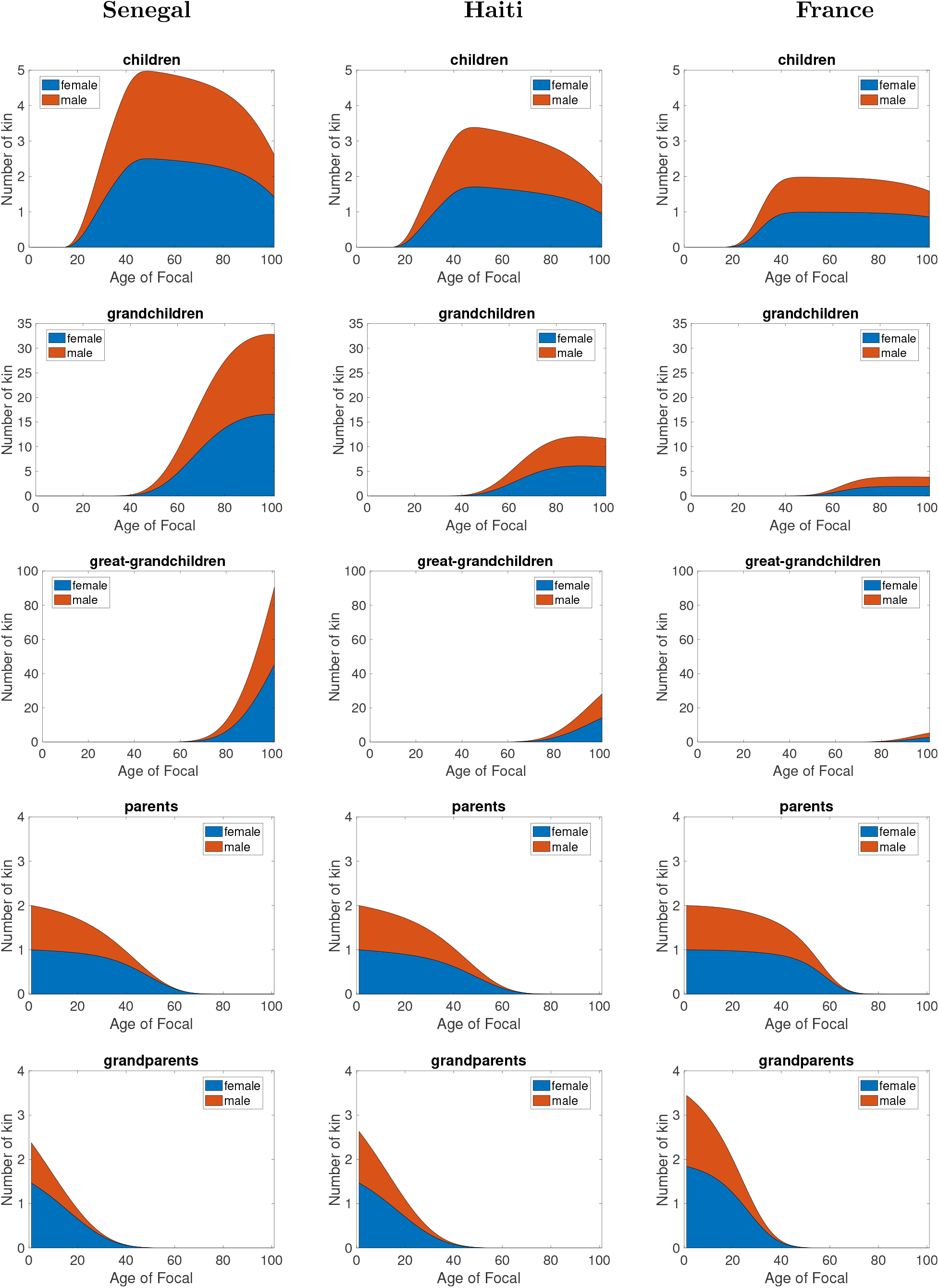
The numbers of male and female kin as a function of the age of Focal. Data for Senegal (left column), Haiti (middle column), and France (right column). For clarity, the *y*-axis are the same across countries, but differ among kin types.

**Figure 8: (part 2 of 2).**
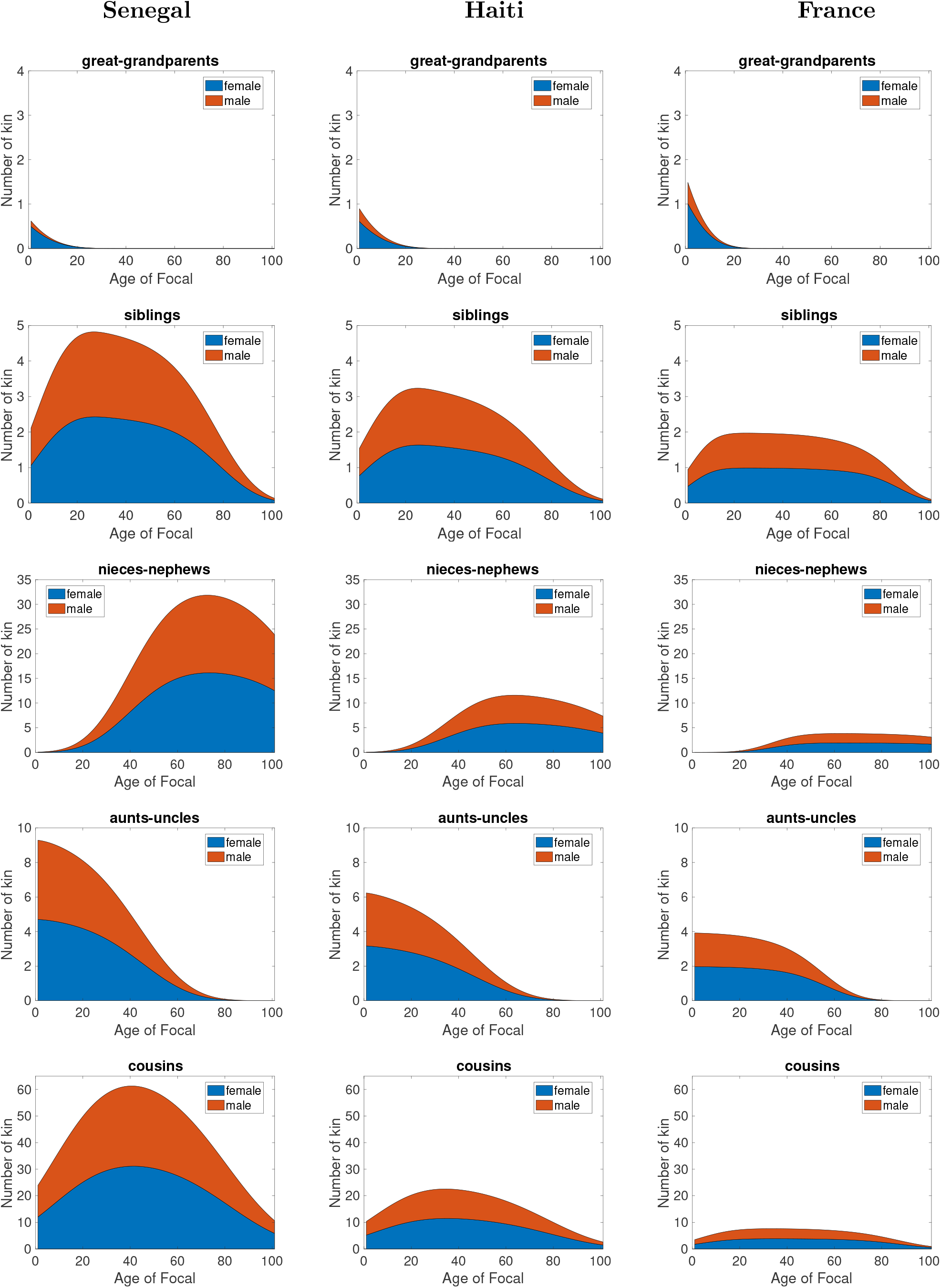
The numbers of male and female kin as a function of the age of Focal. Data for Senegal (left column), Haiti (middle column), and France (right column).

Looking down columns in Figure 8 we see the expected increase in grandchildren over children, and great-grandchildren over grandchildren. The increase is most marked in Senegal, which has the highest fertility. Similarly, we see the expected reduction in grandparents relative to parents, and great-grandparents relative to grandparents. The reduction is most marked in Senegal, with the highest mortality.

The remaining kin types are affected by both fertility and mortality and the patterns are diverse. Niblings compared to siblings behave much like grandchildren when compared to children. Aunts and uncles behave much like parents because by the time of Focal’s birth, new aunts and uncles are not likely to be produced, so the trajectory is dominated by mortality.

Comparing countries across rows in Figure 8, the main differences are those to be expected in comparisons of high to low fertility and high to low mortality populations. The numbers of children, grandchildren, and great-grandchildren decrease from Senegal to France. Parents, grandparents, and great-grandparents decline most rapidly with age in Senegal, and most slowly in France.

Siblings, niblings, aunts-uncles, and cousins are affected, in different ways, by both mortality and fertility schedules. Focal accumulates more of these kin, more rapidly, and loses them more precipitously in Senegal than in Haiti and France.

In general, comparisons across the three countries emphasize the great difference in the sizes and structures of the kinship network created by the differences in mortality and fertility.

### 4.4 Kin sex ratios

What is hinted at, but not clearly apparent, in Figure 8 is the shift in the sex composition of the various kin as Focal ages. Figure 9 shows the sex ratio (number of males divided by number of females), for each type of kin, as a function of the age of Focal.

**Figure 9:**
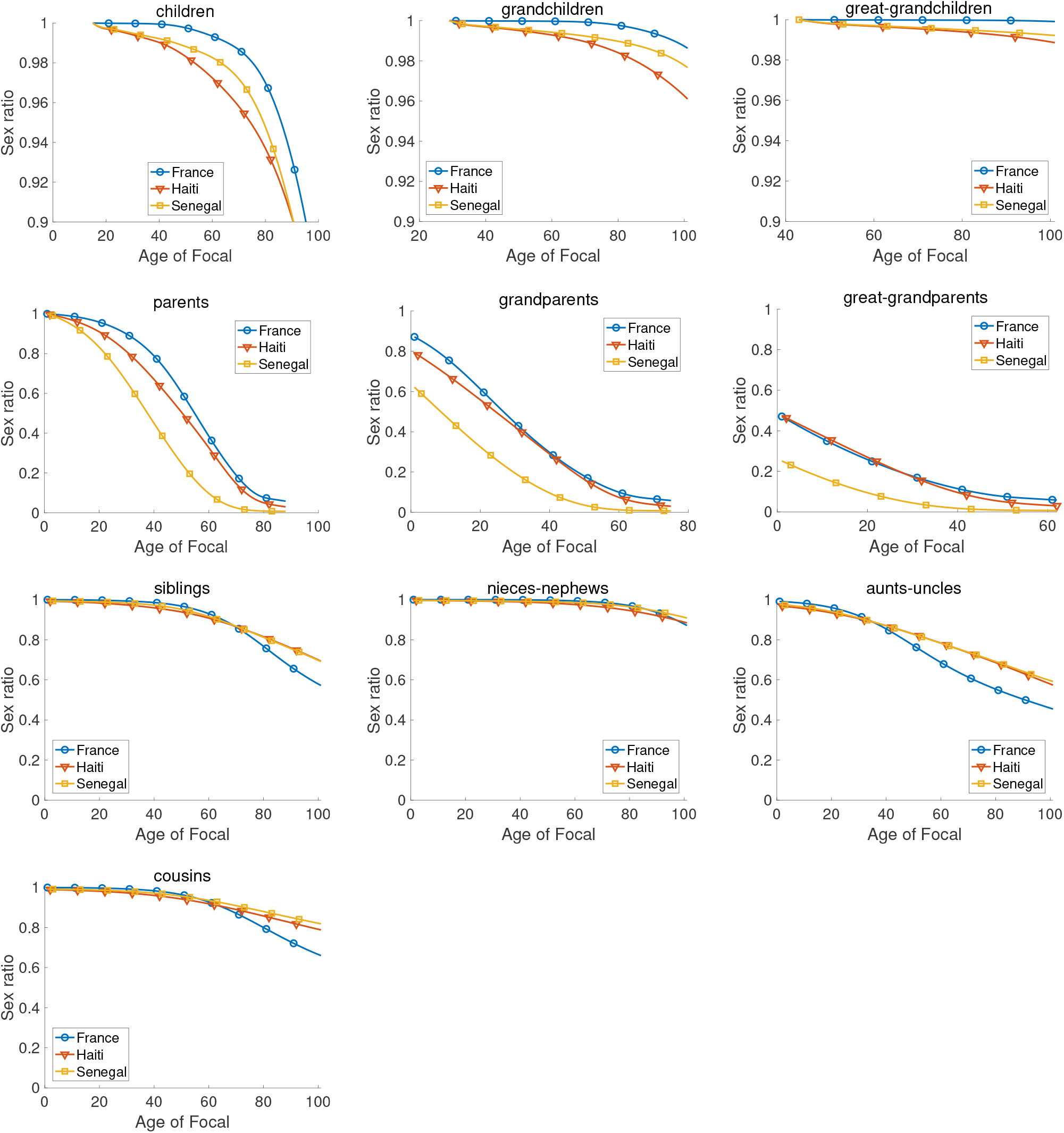
Sex ratios of kin (number of males divided by number of females) as a function of the age of Focal. Note that the *y*-axis scale for children, grandchildren, and great-grandchildren differs from that of other kin, to make patterns visible.

The sex ratio declines (i.e., females come to incrasingly outnumber males) with increasing age of Focal in all cases. Children are of necessity younger than Focal; grandchildren and great-grandchildren even younger. Thus the reduction in sex ratio of her children as Focal ages is small. Parents, grandparents, and great-grandparents are correspondingly older than Focal, and their sex ratios decline dramatically with age of Focal. The declines among siblings (children of Focal’s mother) and cousins (children of Focal’s siblings) are similar. The sex ratios of niblings (the children of Focal’s siblings) are similar to those of the children of Focal herself.

A comparison of the three countries hints at some processes that warrant further investigation. Of the three countries, France shows the largest sex difference in life expectancy (Table 2). Its sex ratio might thus be expected to decline the most rapidly. This is true for some kin (aunts-uncles, cousins) but not for others (children, grandchildren). This suggests that the details of age-specific differences in mortality, rather than overall life expectancy, interact with the age structure of kin to determine the details of the sex ratio patterns. A sensitivity analysis of the kinship model, using methods similar to those of Caswell (2019b) would document the importance of mortality differences at every age to the changes in sex ratios.

## 5 Approximations to the two-sex model

Complete age- and sex-specific mortality and fertility schedules are not always available. In particular, male fertility data are less commonly reported than female fertility data (e.g., Coleman 2000, but see Schoumaker 2019). Fortunately, the structure of the two-sex kinship model readily admits approximations that utilize whatever mortality and fertility data are available. Four such approximations are

**Model 1.** The full two-sex model, as presented here. It utilizes the fertility matrices **F**_f_ and **F**_m_ and the survival matrices **U**_f_ and **U**_m_, see Section 3.

**Model 2.** In the absence of male fertility data, an approximation could use **F**_f_ for both female and male fertility. In this case, set

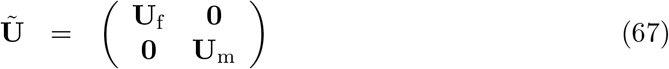

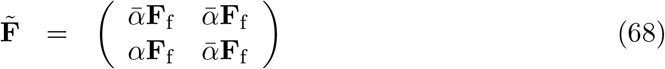

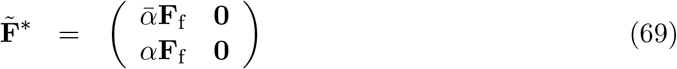

Schoumaker (2019) reports that the difference between male and female fertility is least in countries with low fertility rates; this approximation should, therefore, be most successful in such populations.

**Model 3.** In the absence of male mortality data, an approximation would apply the female mortality schedule to both sexes. In this case, set

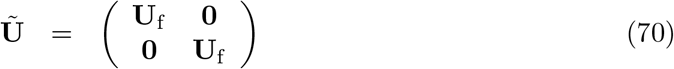

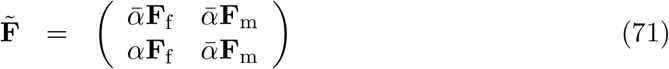

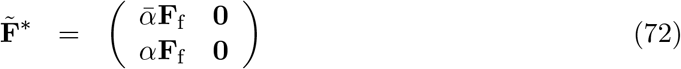

**Model 4.** In the absence of male rates of any kind, males and females could be treated as indistinguishable. I refer to this as the *androgynous approximation*, in which female rates are used for both sexes, leading to

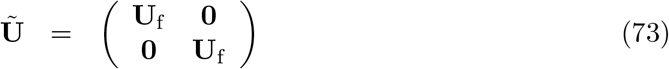

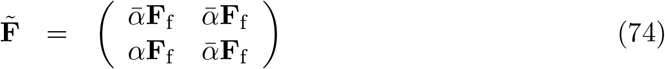

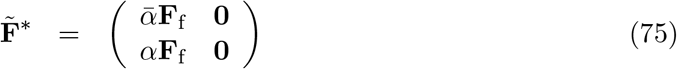

In each of these cases, the calculations proceed as prescribed in Section 3, but using the matrices specified here. The symmetrical cases in which male rates are used in place of missing female rates are an obvious extension.

### 5.1 How good is the androgynous approximation?

The androgynous approximation deserves special consideration because it requires a minimum amount of data. It uses female rates only, treating males and females as identical. However, it does project female and male kin as categories, which can be a useful extension of the one-sex model. A comparison of the full two-sex model and the androgynous approximation reveals the importance of including sex-specific rates. The case of Senegal, with its large differences between male and female fertility, is a worthwhile comparison.

Figure 10 shows the numbers of female and male kin of each type, as a function of the age of Focal, for Model 1 and Model 4, using the rates of Senegal because it has large differences between female and male rates. The differences between the two models are a measure of how much the sex-specific rates contribute to the structure of the kinship network (at least, as measured by numbers of kin).

**Figure 10:**
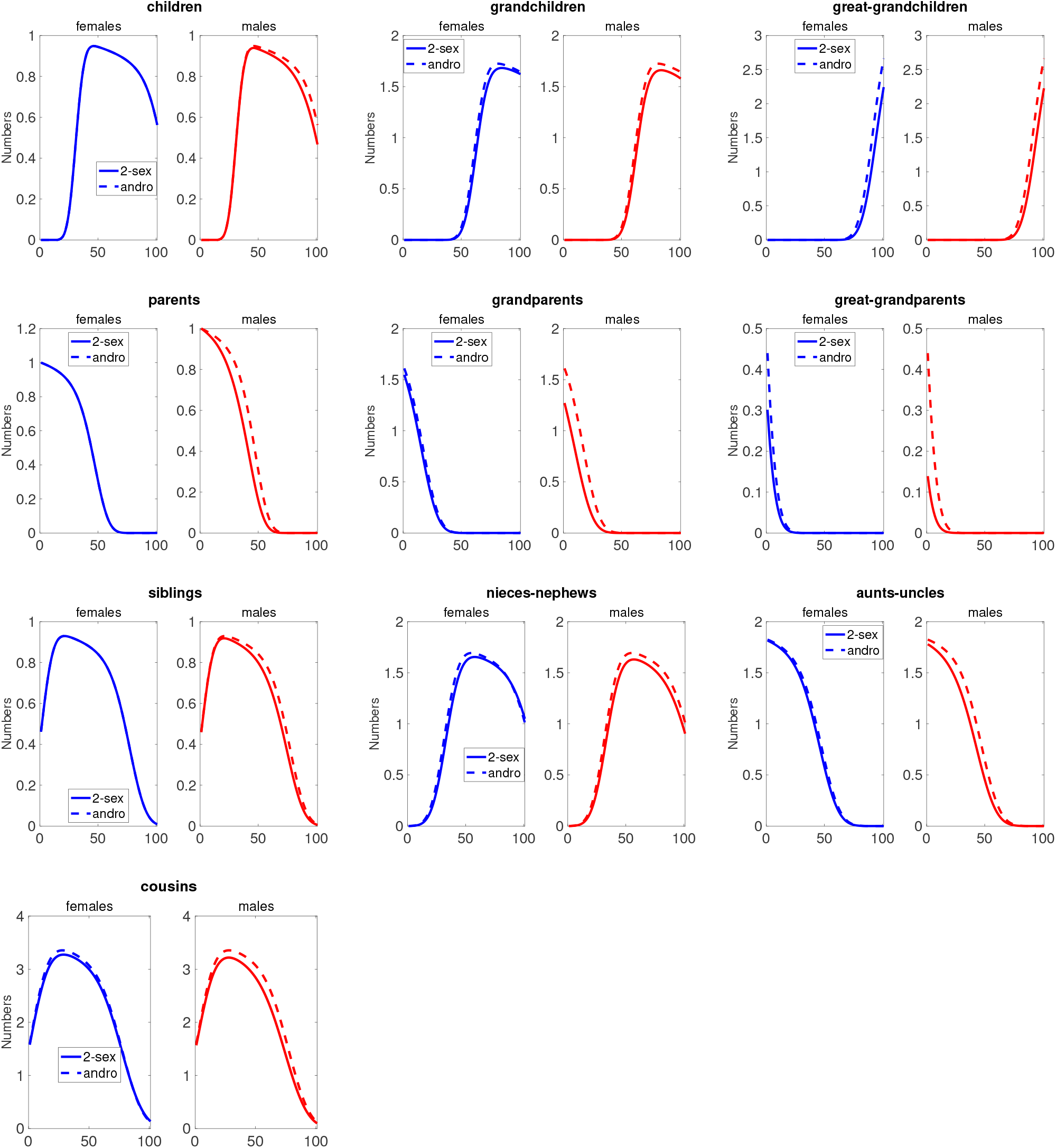
Male and female kin as calculated from the full two-sex Model 1 and the androgynous approximation Model 4. The latter model applies the same (female) rates to both sexes, based on the rates of Senegal.

In general, the differences are small, at most a fraction of an individual. A few kin types (female children, parents, and siblings) show no differences between the two-sex and the androgynous models. The differences between the two models are due to the effects of differences in mortality (two-sex vs. female only) and fertility (two-sex vs. female only). The differences can be decomposed into contributions from these two sources using the Kitagawa-Keyfitz decomposition (Kitagawa, 1955; Keyfitz, 1968, Section 7.4); see Caswell (2010) for description. Figure 11 shows the result of the decomposition for each kin type as a function of the age of Focal.

**Figure 11:**
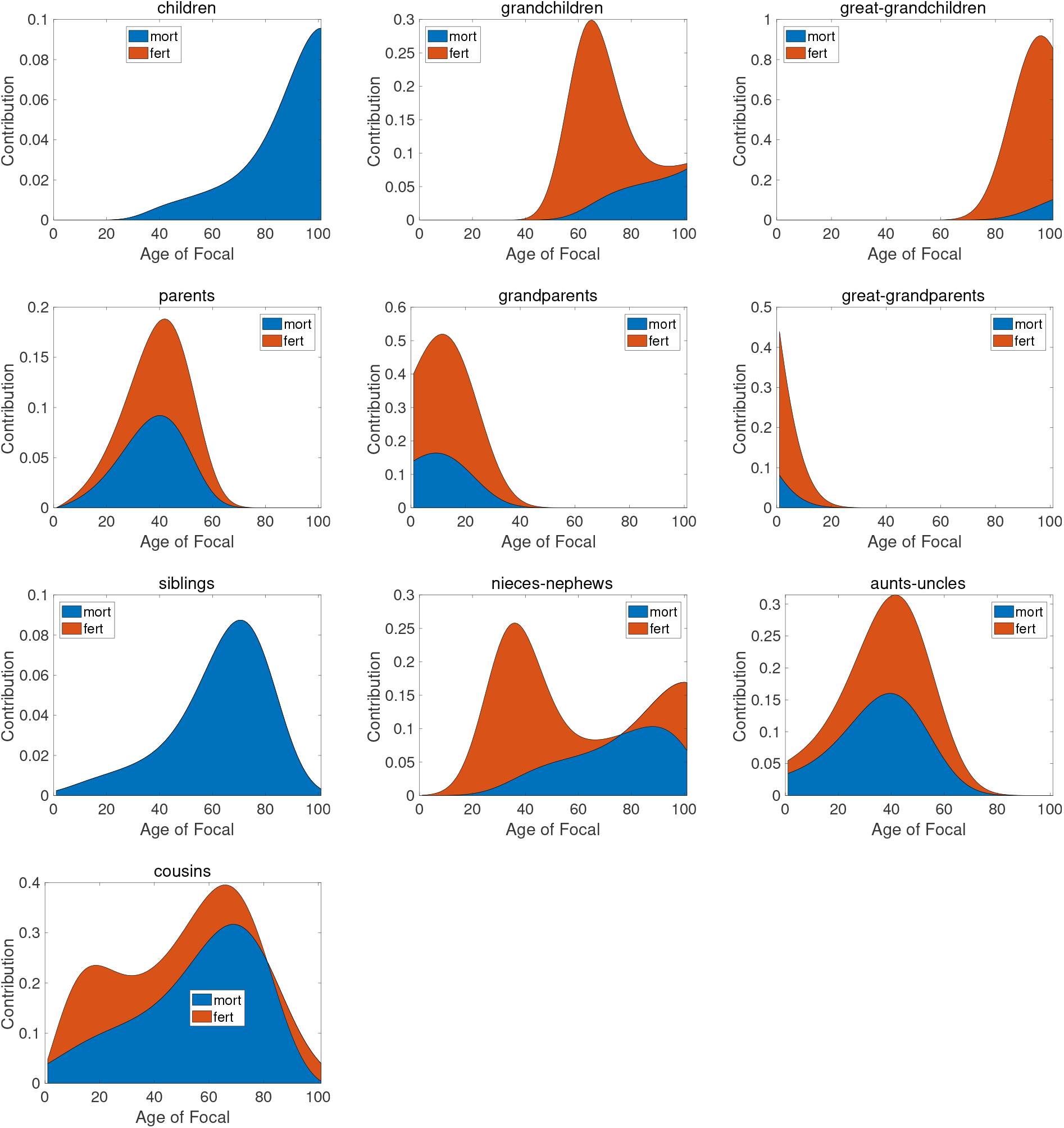
Decomposition analysis. The difference in kin numbers between the andogynous approximation Model 4 and the two-sex Model 1 is decomposed into contributions of differences in mortality and differences in fertility. Based on rates of Senegal.

The sum of the contributions is positive in all cases: the androgynous Model 4 overestimates the kin numbers obtained from the two-sex Model 1. The overall error from the androgynous model is less than one-half of an individual for all kin types except for great-grandchildren at old ages of Focal. Because children and siblings are unaffected by male fertility rates, the differences in kin numbers are totally due to the assumption of identical female and male mortality. All the other kin types are affected by both mortality and fertility assumptions.

#### 5.1.1 The contributions of sex-specific mortality and fertility

#### 5.1.2 Total kin numbers: the GKP factors

In the absence of information on male rates, Goodman, Keyfitz, and Pullum (1974) suggested multiplying the results of a one-sex model by a set of factors (I refer to these as the GKP factors) to obtain kin numbers for a scenario in which female and male rates were identical (i.e., our androgynous approximation Model 4). The GKP factors would multiply daughters by 2, granddaughters by 4, great-granddaughters by 8, mothers by 2, grandmothers by 4, great-grandmothers by 8, sisters by 2, nieces by 4, aunts by 4, and cousins by 8.

Calculations under the Senegal rates show, not surprisingly, that the GKP factors do give exactly the androgynous Model 4 kin numbers. Thus the results of Figure 11) also give the error resulting from using the GKP factors to approximate the results of the full two-sex model.

It should be noted that these comparisons are based on numbers of kin. Other results, especially those concerning sex ratios and sex-specific prevalences, cannot be approximated by one-sex models.

## 6 Discussion

The model presented here analyzes the effects of the sexes by treating sex as an individual state variable in addition to age. As is generally true of such multistate models (Caswell et al., 2018) the form of the model is unchanged ^5^ but the vectors and matrices take on a structure that reflects the enlarged state space:

- the age structure vector **k**(*x*) is replaced by the age×stage vector 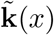,
- the survival matrix **U** is replaced by the block-structured matrix **Ũ**,
- the fertility matrix **F** is replaced with the block structured matrices 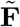 and 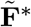, and
- the age at maternity distribution ***π*** is replaced by the maternity and paternity distributions ***π***_f_ and ***π***_m_.

The dynamics of the two-sex model reflect the independence of reproduction by some kin (e.g., sons and daughters of Focal produce grandchildren independently) but dependence in others (e.g., Focal’s mother and father do not produce siblings of Focal independently). The summaries in Table 1 (two-sex) and Tables A-1 (the one-sex model of Caswell 2019a), A-2 (the age×stage-classified model of Caswell 2020), and A-3 (the time-varying model of Caswell and Song 2021) emphasize this formal similarity.

**Table 1:**
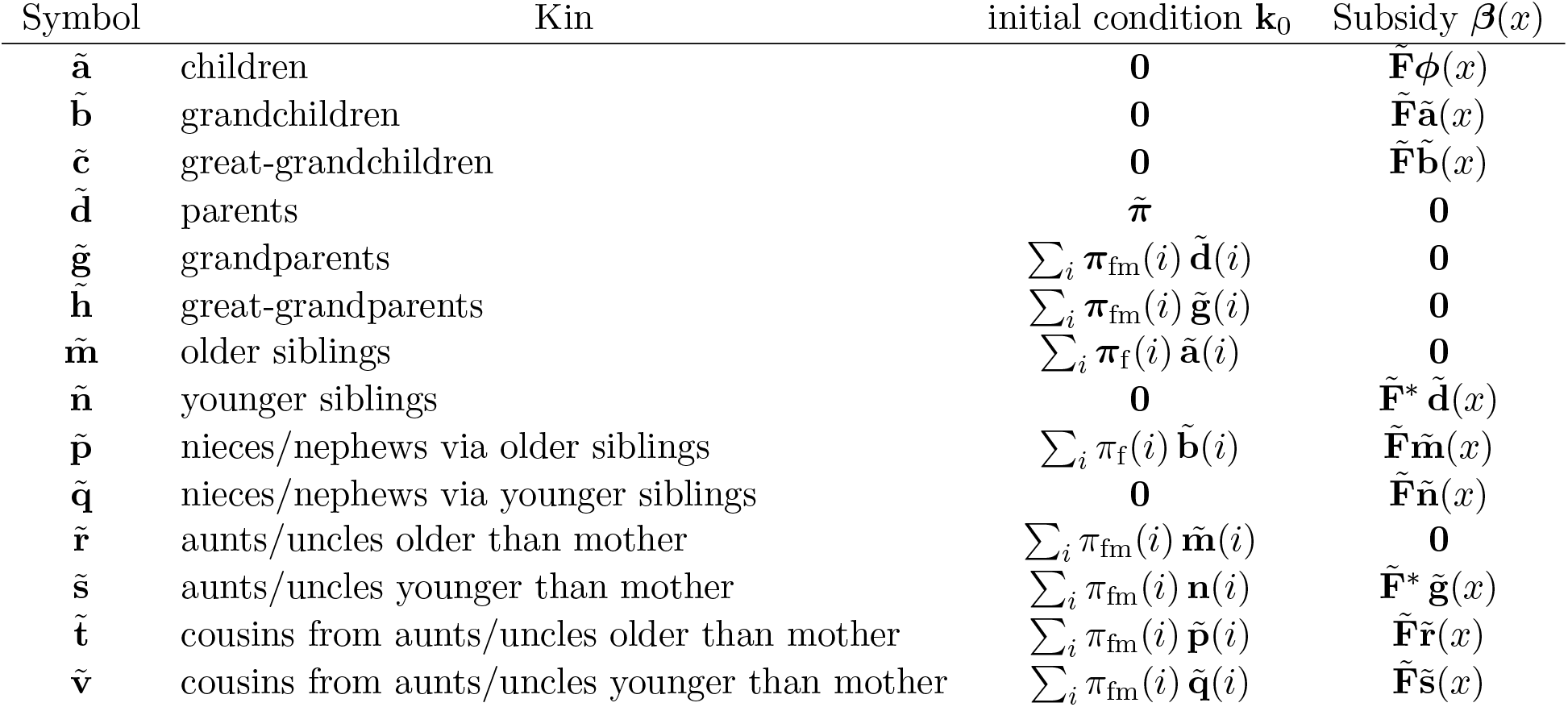
The two-sex kinship model. In this table, ***π***_fm_ = ***π***_f_ + ***π***_m_. Compare this with the age-classified, multistate, and time-varying models in Appendix A.

| Symbol | Kin | initial condition $\mathbf{k}_0$ | Subsidy $\beta(x)$ |
| --- | --- | --- | --- |
| $\tilde{\mathbf{a}}$ | children | $\mathbf{0}$ | $\tilde{\mathbf{F}}\phi(x)$ |
| $\tilde{\mathbf{b}}$ | grandchildren | $\mathbf{0}$ | $\tilde{\mathbf{F}}\tilde{\mathbf{a}}(x)$ |
| $\tilde{\mathbf{c}}$ | great-grandchildren | $\mathbf{0}$ | $\tilde{\mathbf{F}}\tilde{\mathbf{b}}(x)$ |
| $\tilde{\mathbf{d}}$ | parents | $\tilde{\pi}$ | $\mathbf{0}$ |
| $\tilde{\mathbf{g}}$ | grandparents | $\sum_i \pi_{\text{fm}}(i) \tilde{\mathbf{d}}(i)$ | $\mathbf{0}$ |
| $\tilde{\mathbf{h}}$ | great-grandparents | $\sum_i \pi_{\text{fm}}(i) \tilde{\mathbf{g}}(i)$ | $\mathbf{0}$ |
| $\tilde{\mathbf{m}}$ | older siblings | $\sum_i \pi_{\text{f}}(i) \tilde{\mathbf{a}}(i)$ | $\mathbf{0}$ |
| $\tilde{\mathbf{n}}$ | younger siblings | $\mathbf{0}$ | $\tilde{\mathbf{F}}^* \tilde{\mathbf{d}}(x)$ |
| $\tilde{\mathbf{p}}$ | nieces/nephews via older siblings | $\sum_i \pi_{\text{f}}(i) \tilde{\mathbf{b}}(i)$ | $\tilde{\mathbf{F}}\tilde{\mathbf{m}}(x)$ |
| $\tilde{\mathbf{q}}$ | nieces/nephews via younger siblings | $\mathbf{0}$ | $\tilde{\mathbf{F}}\tilde{\mathbf{n}}(x)$ |
| $\tilde{\mathbf{r}}$ | aunts/uncles older than mother | $\sum_i \pi_{\text{fm}}(i) \tilde{\mathbf{m}}(i)$ | $\mathbf{0}$ |
| $\tilde{\mathbf{s}}$ | aunts/uncles younger than mother | $\sum_i \pi_{\text{fm}}(i) \tilde{\mathbf{n}}(i)$ | $\tilde{\mathbf{F}}^* \tilde{\mathbf{g}}(x)$ |
| $\tilde{\mathbf{t}}$ | cousins from aunts/uncles older than mother | $\sum_i \pi_{\text{fm}}(i) \tilde{\mathbf{p}}(i)$ | $\tilde{\mathbf{F}}\tilde{\mathbf{r}}(x)$ |
| $\tilde{\mathbf{v}}$ | cousins from aunts/uncles younger than mother | $\sum_i \pi_{\text{fm}}(i) \tilde{\mathbf{q}}(i)$ | $\tilde{\mathbf{F}}\tilde{\mathbf{s}}(x)$ |

**Table 2:** Male and female life expectancy (*e*_0_), total fertility rate (TFR), and mean ages at maternity and paternity for the data on Senegal, Haiti, and France used in the example calculations. Fertility figures from Schoumaker (2019, Appendix A). Life expectancy data from United Nations (2019).

|  | Senegal | Haiti | France |
| --- | --- | --- | --- |
| $e_0$ female | 67.5 | 63.5 | 85.0 |
| $e_0$ male | 63.8 | 59.3 | 78.7 |
| TFR female | 5.3 | 3.7 | 2.0 |
| TFR male | 11.0 | 5.1 | 2.0 |
| age at maternity | 29.8 | 31.1 | 30.1 |
| age at paternity | 44.1 | 37.0 | 33.5 |

The formal similarity is more than a convenience. It permits the two-sex model to be extended to arbitrary kin types just as the one-sex model has been (although Coste et al. (2021) referred to the method as limited to a specific set of kin, this is not correct). Dead kin and the experience of kin loss can be included just as in Caswell (2019a). Additional state variables can be added as was done for an age×parity model in Caswell (2020). If time series of two-sex rates were available, time variation could be incorporated as in the models of Caswell and Song (2021) and Song and Caswell (2021).

This paper has focused on the numbers and sex ratios of kin. But because the model provides the full age×sex structure for each type of kin, many other kinds of weighted numbers (e.g., dependency ratios) are easily computed. Of particular interest are quantities that might, in general, be called “prevalences” — measures of the occurrence of some property, at specified ages, for each sex. The properties are often medical or health conditions (e.g., Caswell, 2019a, for an analysis of dementia). The prevalences of many conditions are strongly sex dependent. Some cancers, for example, exhibit highly skewed prevalences. In the United States in 2021, new cases of lung cancer, per capita, were 25% higher in men than in women. New cases of kidney cancer were twice as frequent in men, and of bladder cancer four times as frequent (National Cancer Institute, 2021). The prevalence of obstructive sleep apnea in a population in Brazil is both strongly age-dependent (increasing by about three-fold from age 20 to age 80) and sex-dependent (by four-fold controlling for age and other variables) (Tufik et al., 2010).

Sex-specific prevalences, in a more general sense, also appear in non-medical conditions. For example, the income level of kin could be an important issue for Focal, and the gender gap in pay rates is widespread and well known. In the EU as of 2014, women earned from 1% (Romania) to 24% (Estonia) less than men, with an average gap of 14% (Boll and Lagerman, 2018). Song and Caswell (2021) analyzed unemployment among kin, but had to rely on applying the GKP factors to a one-sex model, and thus could not incorporate sex-specific unemployment. Such an analysis would be possible using the two-sex model.

Many gender-specific demographic and sociological variables can be related to kinship structures using this model. In light of this goal, the apparent adequacy of the andogynous approximation to the full two-sex model, in a case (Senegal) with quite large male-female differences, is encouraging. It appears safe to use the approximations listed in Section 5, especially in cases where both male and female mortality are available, but male fertility is lacking. Notice that the differences in numbers of kin between the two-sex model and the androgynous approximation (Figures 10 and 11) are much smaller than the differences in kin numbers among the three countries examined here (Figure 8). More comparative research is warranted.

The possible use of model fertility schedules (e.g., Coale and Trussell, 1974) when measured fertility data are lacking deserves further research. Paget and Timaeus (1994), extending work of Booth (1984), describe a relational model for male fertility,^6^ The relational model produces fertility patterns (their Figure 2) not unlike those in Figure 7.

Despite the growing list of factors included in the matrix kinship models, limitations and open research questions remain. Multiple births, for example, are not included. This is easily addressed and is especially important in biodemography and population biology if one is interested in the kinship structure of species whose fertility patterns are very different from ours (Caswell, 2021).

The distributions of ages at maternity and paternity of Focal’s mother and father determine initial conditions for kin. These ages have been treated here as independent; an obvious extension would be to use a joint distribution reflecting the age distribution of couples.

This model, like previous models, is limited to consanguineal, biological kin. The challenge of incorporating affinal kin, stepkin, kin by marriage, and blended families is an open research question. Perhaps a first step towards a solution would be to combine the kinship networks of two Focal individuals of different ages to provide a picture of a blended set of kin.^7^ The ages of the Focal individuals to be combined might be chosen from a distribution of the ages of parents forming blended families.

Finally, it is important to recall that the projections of the kin populations provide mean age-sex structures, over the distributions produced by the survival and fertility probabilities. Because the kin populations are small, there will be an (as yet unknown) degree of stochastic variation around those means.

In summary, the matrix theoretic model makes it possible to expand the demographic detailed included the analysis of a kinship network: from age distributions to multistate age×stage distributions, from time-invariant to time-varying demographic rates, and now from one-sex to two-sex models. It has resulted in an increasingly rich set of demographic outcomes, including many of great demographic interest, including (but not limited to) bereavement, dependency, prevalences, family sizes, and sex ratios. All these are now subject to analysis in terms of demographic rates in a kinship setting.

## 7 Acknowledgments

This research was supported by the European Research Council under the European Union’s Horizon 2020 research and innovation program through ERC Advanced Grant 788195. Motion Coffee in Amsterdam provided a pleasant environment for work on this and other projects. I thank Xi Song for discussions, and am especially grateful to Bruno Schoumaker for providing the male fertility data used in the examples here.

## A One-sex, time-varying, and multistate kinship models

This appendix presents tables showing the expressions for the kinship dynamics in one-sex, multistate, and time-varying kinship models. Comparison of these tables with with Table 1 shows the formal similarity of the four models. Tables are modified from their cited sources under the terms of a Creative Commons Attribution license.

**Table A-1:** The age-classified, time-invariant, one-sex kinship model of Caswell (2019a).

| Symbol | Kin | initial condition $\mathbf{k}_0$ | Subsidy $\beta(x)$ |
| --- | --- | --- | --- |
| **a** | daughters | **0** | **F**$\mathbf{e}_x$ |
| **b** | granddaughters | **0** | **F**$\mathbf{a}(x)$ |
| **c** | great-granddaughters | **0** | **F**$\mathbf{b}(x)$ |
| **d** | mothers | $\boldsymbol{\pi}$ | **0** |
| **g** | grandmothers | $\sum_i \pi_i \mathbf{d}(i)$ | **0** |
| **h** | great-grandmothers | $\sum_i \pi_i \mathbf{g}(i)$ | **0** |
| **m** | older sisters | $\sum_i \pi_i \mathbf{a}(i)$ | **0** |
| **n** | younger sisters | **0** | **F**$\mathbf{d}(x)$ |
| **p** | nieces via older sisters | $\sum_i \pi_i \mathbf{b}(i)$ | **F**$\mathbf{m}(x)$ |
| **q** | nieces via younger sisters | **0** | **F**$\mathbf{n}(x)$ |
| **r** | aunts older than mother | $\sum_i \pi_i \mathbf{m}(i)$ | **0** |
| **s** | aunts younger than mother | $\sum_i \pi_i \mathbf{n}(i)$ | **F**$\mathbf{g}(x)$ |
| **t** | cousins from aunts older than mother | $\sum_i \pi_i \mathbf{p}(i)$ | **F**$\mathbf{r}(x)$ |
| **v** | cousins from aunts younger than mother | $\sum_i \pi_i \mathbf{q}(i)$ | **F**$\mathbf{s}(x)$ |

**Table A-2:** Summary of the age×stage-classified kinship model of Caswell (2020). Matrices and vectors bearing tildes (e.g., **ã**) age×stage block-structured.

| Symbol | Kin | initial condition $\mathbf{k}_0$ | Subsidy $\beta(x)$ |
| --- | --- | --- | --- |
| $\tilde{\phi}$ | Focal | $\tilde{\phi}_0$ | **0** |
| $\tilde{\mathbf{a}}$ | daughters | **0** | $\tilde{\mathbf{F}}\tilde{\phi}(x)$ |
| $\tilde{\mathbf{b}}$ | granddaughters | **0** | $\tilde{\mathbf{F}}\tilde{\mathbf{a}}(x)$ |
| $\tilde{\mathbf{c}}$ | great-granddaughters | **0** | $\tilde{\mathbf{F}}\tilde{\mathbf{b}}(x)$ |
| $\tilde{\mathbf{d}}$ | mothers | $\tilde{\boldsymbol{\pi}}$ | **0** |
| $\tilde{\mathbf{g}}$ | grandmothers | $\sum_i \pi_i^{\text{age}} \tilde{\mathbf{d}}(i)$ | **0** |
| $\tilde{\mathbf{h}}$ | great-grandmothers | $\sum_i \pi_i^{\text{age}} \tilde{\mathbf{g}}(i)$ | **0** |
| $\tilde{\mathbf{m}}$ | older sisters | $\sum_i \pi_i^{\text{age}} \tilde{\mathbf{a}}(i)$ | **0** |
| $\tilde{\mathbf{n}}$ | younger sisters | **0** | $\tilde{\mathbf{F}}\tilde{\mathbf{d}}(i)$ |
| $\tilde{\mathbf{p}}$ | nieces via older sisters | $\sum_i \pi_i^{\text{age}} \tilde{\mathbf{b}}(i)$ | $\tilde{\mathbf{F}}\tilde{\mathbf{m}}(x)$ |
| $\tilde{\mathbf{q}}$ | nieces via younger sisters | **0** | $\tilde{\mathbf{F}}\tilde{\mathbf{n}}(i)$ |
| $\tilde{\mathbf{r}}$ | aunts older than mother | $\sum_i \pi_i^{\text{age}} \tilde{\mathbf{m}}(i)$ | **0** |
| $\tilde{\mathbf{s}}$ | aunts younger than mother | $\sum_i \pi_i^{\text{age}} \tilde{\mathbf{n}}(i)$ | $\tilde{\mathbf{F}}\tilde{\mathbf{g}}(x)$ |
| $\tilde{\mathbf{t}}$ | cousins: aunts older than mother | $\sum_i \pi_i^{\text{age}} \tilde{\mathbf{p}}(i)$ | $\tilde{\mathbf{F}}\tilde{\mathbf{r}}(x)$ |
| $\tilde{\mathbf{v}}$ | cousins: aunts younger than mother | $\sum_i \pi_i^{\text{age}} \tilde{\mathbf{q}}(i)$ | $\tilde{\mathbf{F}}\tilde{\mathbf{s}}(x)$ |

**Table A-3:** The time-varying kinship model of Caswell and Song (2021). Age (of Focal) and time are denoted by *x* and *t*, respectively. For each type of kin, the relevant age boundary condition, survival dynamics, and reproductive subsidy are shown.

| Kin | age boundary | survival | subsidy $\beta(x)$ |
| --- | --- | --- | --- |
| $\mathbf{a}(x, t)$ daughters | $\mathbf{0}$ | $\mathbf{U}_t \mathbf{a}(x, t)$ | $\mathbf{F}_t \mathbf{e}_x$ |
| $\mathbf{b}(x, t)$ granddaughters | $\mathbf{0}$ | $\mathbf{U}_t \mathbf{b}(x, t)$ | $\mathbf{F}_t \mathbf{a}(x, t)$ |
| $\mathbf{c}(x, t)$ great-granddaughters | $\mathbf{0}$ | $\mathbf{U}_t \mathbf{c}(x, t)$ | $\mathbf{F}_t \mathbf{b}(x, t)$ |
| $\mathbf{d}(x, t)$ mothers | $\boldsymbol{\pi}(t)$ | $\mathbf{U}_t \mathbf{d}(x, t)$ | $\mathbf{0}$ |
| $\mathbf{g}(x, t)$ grandmothers | $\sum_i \pi_i(t) \mathbf{d}(i, t)$ | $\mathbf{U}_t \mathbf{g}(x, t)$ | $\mathbf{0}$ |
| $\mathbf{h}(x, t)$ great-grandmothers | $\sum_i \pi_i(t) \mathbf{g}(i, t)$ | $\mathbf{U}_t \mathbf{h}(x, t)$ | $\mathbf{0}$ |
| $\mathbf{m}(x, t)$ older sisters | $\sum_i \pi_i(t) \mathbf{a}(i, t)$ | $\mathbf{U}_t \mathbf{m}(x, t)$ | $\mathbf{0}$ |
| $\mathbf{n}(x, t)$ younger sisters | $\mathbf{0}$ | $\mathbf{U}_t \mathbf{n}(x, t)$ | $\mathbf{F}_t \mathbf{d}(x, t)$ |
| $\mathbf{p}(x, t)$ nieces via older sisters | $\sum_i \pi_i(t) \mathbf{b}(i, t)$ | $\mathbf{U}_t \mathbf{p}(x, t)$ | $\mathbf{F}_t \mathbf{m}(x, t)$ |
| $\mathbf{q}(x, t)$ nieces via younger sisters | $\mathbf{0}$ | $\mathbf{U}_t \mathbf{q}(x, t)$ | $\mathbf{F}_t \mathbf{n}(x, t)$ |
| $\mathbf{r}(x, t)$ aunts older than mother | $\sum_i \pi_i(t) \mathbf{m}(i, t)$ | $\mathbf{U}_t \mathbf{r}(x, t)$ | $\mathbf{0}$ |
| $\mathbf{s}(x, t)$ aunts younger than mother | $\sum_i \pi_i(t) \mathbf{n}(i, t)$ | $\mathbf{U}_t \mathbf{s}(x, t)$ | $\mathbf{F}_t \mathbf{g}(x, t)$ |
| $\mathbf{t}(x, t)$ cousins; aunts older than mother | $\sum_i \pi_i(t) \mathbf{p}(i, t)$ | $\mathbf{U}_t \mathbf{t}(x, t)$ | $\mathbf{F}_t \mathbf{r}(x, t)$ |
| $\mathbf{v}(x, t)$ cousins; aunts younger than mother | $\sum_i \pi_i(t) \mathbf{q}(i, t)$ | $\mathbf{U}_t \mathbf{v}(x, t)$ | $\mathbf{F}_t \mathbf{s}(x, t)$ |

## Footnotes

1 An extreme case is that of Moulay Ismael the Bloodthirsty, the Emperor of Morocco (1672–1727). This unpleasant individual is reported to have sired 888 children. A recent analysis by Oberzaucher and Grammer (2014) has concluded that it is indeed possible to that he could have done so.

2 There exist situations in which the full structure of Figure 2 and equation (7) would be of interest. For example, evolutionary calculations based on kin selection depend on the sharing of genes between individuals. Because maternity is more certain than paternity, Focal is more certain that she shares 1/4 of her genes with the children of her daughters, 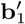 and 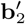 than she is for the children of her sons, 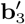 and 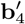. In general, each passage through a male descendant will introduce some uncertainty into the inheritance. A related issue arises with respect to mitochondrial DNA, which is passed from mothers to offspring, and Y chromosomes, which are transmitted from fathers to sons; calculations involving these forms of inheritance might benefit from being able to distinguish kin by lines of descent. See Tanskanen and Danielsbacka (2019), especially their Table 2.2, for a summary of these issues.

3 A slight modification of the initial conditions for older and younger siblings can account for the possibility of multiple births.

4 https://population.un.org/wpp/Download/Archive/Standard/

5 This invariance of form, and hence of analysis, is part of the definition of “formal” in formal demography.

6 I must confess that I do not understand this paper.

7 Much as in the case of: 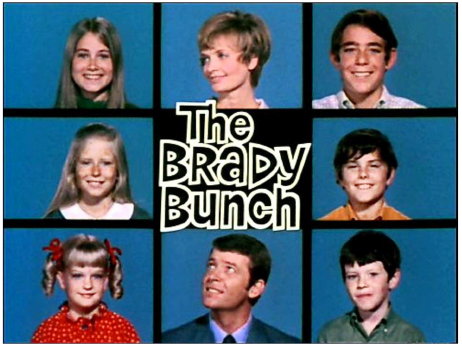

